# Stromal Hippo-YAP signaling in stem cell niche controls intestinal homeostasis

**DOI:** 10.1101/2022.05.12.491640

**Authors:** Kyvan Dang, Alka Singh, Jennifer L. Cotton, Zhipeng Tao, Haibo Liu, Lihua J. Zhu, Xu Wu, Junhao Mao

## Abstract

Intestinal homeostasis is tightly regulated by the reciprocal interaction between gut epithelium and adjacent mesenchyme. The mammalian Hippo-YAP pathway is intimately associated with intestinal epithelial homeostasis and re-generation; however, its role in postnatal gut mesenchyme remains poorly defined. We find that, although removal of the core Hippo kinases Lats1/2 or activation of YAP in adult intestinal smooth muscle has largely no effect; Hippo-YAP signaling in Gli1/PDGFR-expressing intestinal stromal cells is critical to maintain the stem cell niche. We show that YAP/TAZ activation drives over-proliferation and suppresses smooth muscle actin expression in the niche-forming Gli1^+^ mesenchymal progenitors. In addition, mesenchymal YAP/TAZ activation disrupts the epithelial-mesenchymal crosstalk by promoting Wnt ligand production, leading to epithelial Wnt pathway activation. Our data also reveal that YAP/TAZ are upregulated in the stroma during DSS-induced injury and stromal YAP activation promotes intestinal epithelial regeneration. Altogether, our data identify an essential requirement for stromal Hippo-YAP signaling in the stem cell niche during intestinal homeostasis.

**HIGHTLIGHTS:**

- Lats1/2 control proliferation and differentiation of adult gut mesenchymal progenitors.
- Mesenchymal YAP/TAZ promotes Wnt ligand production in intestinal stem cell niche.
- YAP/TAZ is up-regulated in the mesenchyme during intestinal injury.
- Stromal YAP activation promotes intestinal epithelial regeneration.

## INTRODUCTION

The mammalian gastrointestinal (GI) tract is composed of the endodermis-derived epithelium surrounded by adjacent mesoderm-derived mesenchyme. Continuous cross-talks between these two tissue layers are critical for intestinal development and postnatal homeostasis (Kedinger et al., 1998). However, in comparison to the intestinal epithelium, the genetic and molecular mechanisms governing mesenchymal growth, differentiation, and homeostasis are under-studied and poorly understood. It is known that mesenchymal/stromal dysfunction is closely associated with human digestive tract diseases (Kedinger et al., 1998). Recent studies have also demonstrated that subepithelial mesenchymal cells form the essential Wnt-secreting niche needed to sustain intestinal epithelial stem cells in the crypt (Degirmenci et al., 2018; Greicius et al., 2018; Shoshkes-Carmel et al., 2018; Valenta et al., 2016), therefore playing a pivotal role in controlling intestinal homeostasis.

The Hippo pathway, originally identified in *Drosophila* as an organ size control pathway, consists of a core kinase cascade, Mst1/2 and Lats1/2, which phosphorylate and inactivate the transcriptional co-activators YAP and TAZ. Upon upstream Hippo signaling inactivation, YAP/TAZ translocate into the nucleus and interact with the TEAD family of transcription factors, thereby inducing downstream gene expression (Gregorieff and Wrana, 2017; Halder and Camargo, 2013; Hong et al., 2016; Zanconato et al., 2016; Zheng and Pan, 2019). Works from us and many others demonstrated a critical requirement for epithelial Hippo signaling in intestinal stem cell maintenance and regeneration (Barry et al., 2013; Cai et al., 2010; Cheung et al., 2020; Gregorieff et al., 2015; Gregorieff and Wrana, 2017; Li et al., 2020; Yui et al., 2018). Furthermore, our recent work also uncovered the previously unappreciated requirement of YAP/TAZ in the embryonic gut mesenchyme, linking the Hippo-YAP pathway to intestinal mesenchymal homeostasis (Cotton et al., 2017). We found that YAP/TAZ function as a molecular switch to coordinate mesenchymal growth and patterning in the developing gut (Cotton et al., 2017). However, the precise role of Hippo-YAP signaling in the stromal compartment of adult intestine remains largely unknown.

In this study, we explored the function of stromal Lats1/2, the core Hippo kinases, and YAP during intestinal homeostasis, focusing on their requirement in the mesenchymal progenitor populations within the stem cell niche. We found that Hippo signaling controls proliferation and differentiation of the Gli1^+^ mesenchymal cells and regulates Wnt ligand production in the stem cell niche. Our data show that YAP activation in the mesenchymal cells promotes intestinal epithelial regeneration following injury.

## RESULTS

### Disruption of Hippo-YAP signaling in intestinal gut smooth muscle

To explore mesenchymal Hippo-YAP signaling during intestinal homeostasis, we first examined the expression of YAP/TAZ in adult intestine. Immunohistochemical (IHC) staining and immunoblotting for YAP/TAZ showed that YAP/TAZ were highly expressed in the intestinal mesenchyme in comparison to the intestinal epithelium (Figure 1A, B). Interestingly, we found that the YAP/TAZ levels were significantly higher in the outer smooth muscle layers than those in the subepithelial stromal cells (Figure 1B), and it is consistent with our quantitative RT-PCR (qPCR) analysis of the transcription of *bona fide* YAP target genes, *Ctgf* and *Ankrd1*, in these cells (Figure 1C).

**Figure 1.**
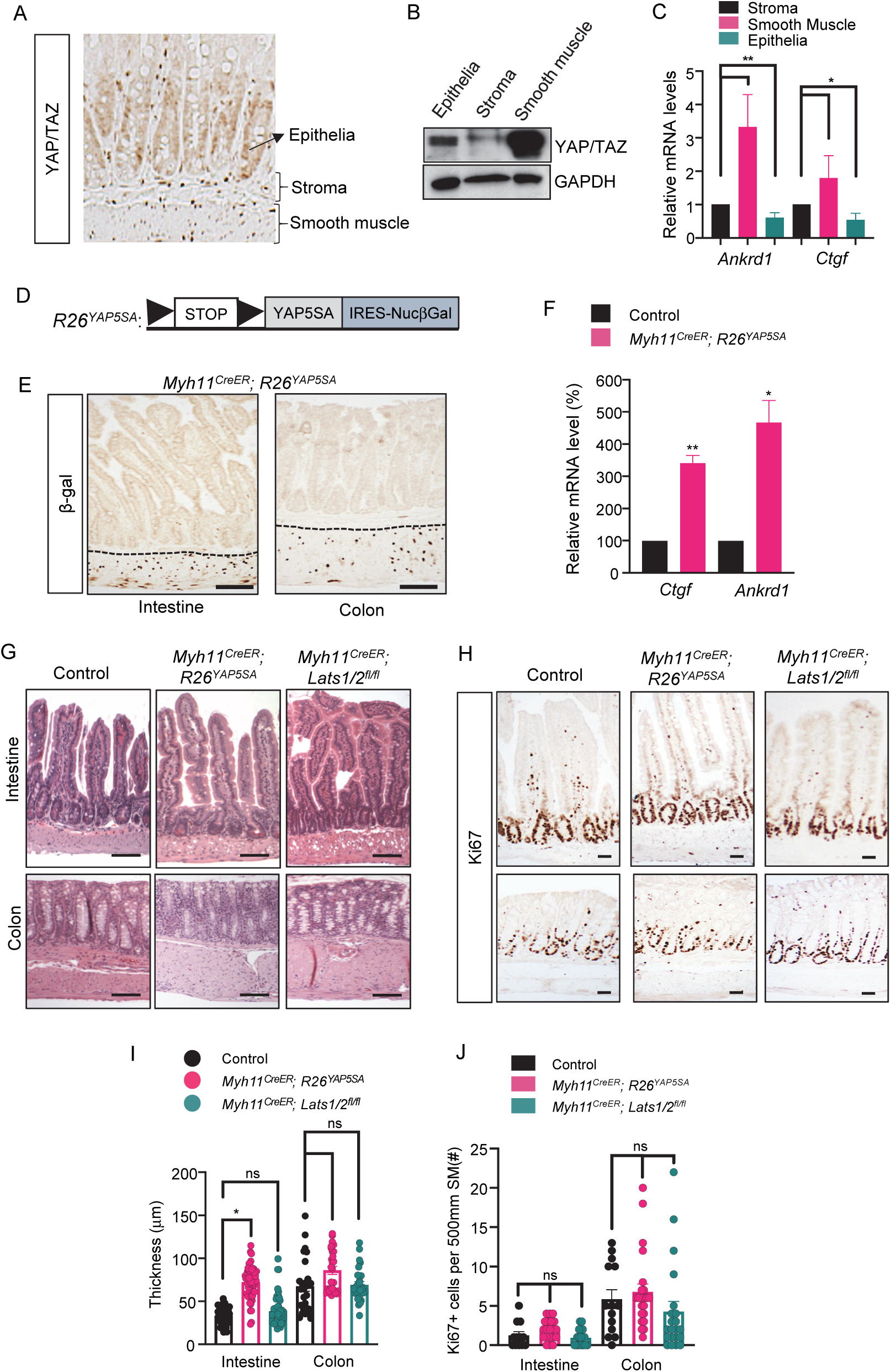
Disruption of Hippo-YAP signaling in intestinal smooth muscle. Immunohistochemistry of YAP/TAZ in control intestine, showing YAP/TAZ staining in intestinal epithelia, subepithelial stroma and smooth muscle layers. (**B**) Immunoblot analysis of YAP/TAZ and GAPDH in intestinal epithelia, stroma and smooth muscle. (**C**) Real-time PCR analysis showing relative mRNA levels of *Ankrd1* and *Ctgf* mRNA levels in intestinal stroma, smooth muscle, and epithelia. (n=3 biological replicate, n=3 technical replicate). (**D**) Schematic representation of the *R26*^*YAP5SA*^ allele. (**E**) Immunohistochemistry of β-galactosidase (β-gal) in intestine and colon of *Myh11*^*CreER*^; *R26*^*YAP5SA*^ animals. Stroma-smooth muscle boundary is indicated by dashed line. Scale bar, 50 μm. (**F**) Real-time PCR analysis of *Ctgf* and *Ankrd1* mRNA levels in control and *Myh11*^*CreER*^; *R26*^*YAP5SA*^ mutant smooth muscle cells. (n=3 biological replicate, n=3 technical replicate). (**G**) Histology of intestine and colon of control, *Myh11*^*CreER*^; *R26*^*YAP5SA*^, and *Myh11*^*CreER*^; *Lats1/2*^*fl/fl*^ animals. Scale bar, 50 μm. (**H**) Immunohistochemical Ki67 staining in intestine and colon of control, *Myh11*^*CreER*^; *R26*^*YAP5SA*^, and *Myh11*^*CreER*^; *Lats1/2*^*fl/fl*^ animals. Scale bar, 50 μm. (**I**) Quantification of thickness of smooth muscle layer in the intestine and colon of control, *Myh11*^*CreER*^; *R26*^*YAP5SA*^, and *Myh11*^*CreER*^; *Lats1/2*^*fl/fl*^ animals. n=27 per group per anatomical location. (n= 3 animals per group, >27 measurements per group) (**J**) Quantification of Ki67^+^ smooth muscle cells in the smooth muscle layer in intestine and colon of WT, *Myh11*^*CreER*^; *R26*^*YAP5SA*^, and *Myh11*^*CreER*^; *Lats1/2*^*fl/fl*^ animals. (n= 2 animals per group, 12 fields per group) Data are mean ± SEM., * p<0.05, ** p<0.001, ns: not significant. See also Figure S1.

To further investigate the role of Hippo-YAP signaling in smooth muscle cells, we either crossed the conditional active YAP allele, *R26*^*YAP5SA*^ (Cotton et al., 2017) or the floxed *Lats1/2* alleles (*Lats1/2*^*fl/fl*^) (Yi et al., 2016), to smooth muscle cell-specific *Myh11*^*CreER*^ (Wirth et al., 2008). Recombination was induced in animals via Tamoxifen (TM) intraperitoneal (I.P.) injections at post-natal day 30 (P30). In the *R26*^*YAP5SA*^ allele we recently generated, YAP5SA, a constitutively active form of YAP that has five canonical LATS phosphorylation sites mutated from serine to alanine (Zhao et al., 2007) and carries a C-terminal IRES-nuclear β-galactosidase tag, was targeted into the *Rosa26* locus (Cotton et al., 2017) (Figure 1D). Immunohistochemical (IHC) analysis of β-galactosidase in *Myh11*^*CreER*^; *R26*^*YAP5SA*^ mutants following Tamoxifen injection showed nuclear signal exclusively in the smooth muscles of both intestine and colon, indicating transgenic YAP expression (Figure 1E). Elevated expression levels of YAP targets *Ctgf* and *Cyr61* via qRT-PCR of mutant smooth muscle cells compared to control further confirmed YAP activation (Figure 1F).

Despite Lats1/2 deletion or constitutively active YAP expression in the smooth muscle layers, overall intestinal morphology remained largely unchanged, even 80 days following Tamoxifen induction in both *Myh11*^*CreER*^; *R26*^*YAP5SA*^ and *Myh11*^*CreER*^; *Lats1/2*^*fl/fl*^ mutants (Figure 1G). We found that Lats1/2 deletion or YAP activation in gut smooth muscle cells could not induce proliferation (Figure 1H, J), and there is little or no change in smooth muscle thickness between control and mutant animals (Figure 1I). In addition, we did not observe significant change in growth and differentiation of the intestinal epithelia in mutant animals (Figure S1). Although we could not observe mutant animals beyond 90 days after Tamoxifen injection due to overall health deterioration of mutant mice, including severe skin lesions, these data suggested that disruption of Hippo-YAP signaling via Lats1/2 deletion or constitutive YAP activation by *Myh11*^*CreER*^ did not generate an obvious phenotype in the terminally differentiated intestinal smooth muscle cells.

### Lats1/2 deletion and YAP activation drive intestinal mesenchymal overgrowth

We next explored Hippo-Yap function in the subepithelial compartment of the intestinal mesenchyme. Our effort was focused on the critical mesenchymal populations that form the intestinal stem cell niche. Gli1-expressing cells has recently been identified as the mesenchymal progenitor population serving as a major source of Wnt ligand production that is critical for the maintenance of the stem cell niche (Degirmenci et al., 2018). PDGFRs are also known to be expressed in the pericryptal myofibroblasts that secrete Wnt ligands and agonists for intestinal stem cells (Greicius et al., 2018). We performed immunofluorescence staining of PDGFRβ in the intestine and found that, similar to Gli1, PDGFRβ was also expressed in a population of pericryptal mesenchymal cells including myofibroblasts expressing α-smooth muscle actin (SMA) (Figure S2). This data is consistent with the previous study showing co-expression of Gli1 and PDGFRβ in a subset of subepithelial stromal cells in the stem cell niche (Degirmenci et al., 2018).

To determine the role of Hippo-YAP signaling in these stromal cell populations, we crossed the *Gli1*^*CreER*^ (Ahn and Joyner, 2005) and *Pdgfrb*^*CreER*^ (Gerl et al., 2015) knock-in inducible Cre mice to the floxed *Lats1/2* alleles or *R26*^*YAP5SA*^. The mutant mice were treated with Tamoxifen via IP injection at P30 to induce Cre-mediated recombination. At two months following Tamoxifen induction, pronounced overgrowths were detected in the small intestine and colon of the mutant animals (Figure 2A, Figure S3). These overgrowth areas exhibited disturbed epithelial architecture and drastic mesenchymal expansion (Figure 2A, Figure S3). In the *Gli1*^*CreER*^; *Lats1/2*^*fl/fl*^ mice, IHC of YAP and TAZ showed nuclear YAP/TAZ staining within the expanded mesenchyme (Figure 2B), indicating YAP/TAZ activation induced by Lats1/2 deletion. In *Gli1*^*CreER*^; *R26*^*YAP5SA*^ and *Pdgfrb*^*CreER*^; *R26*^*YAP5SA*^ mice, IHC of nuclear β-galactosidase demonstrated that YAP5SA transgene expression in the overgrowth regions occurred exclusively in the mesenchymal cells (Figure 2C, D, Supplemental Figure 3), which is confirmed by Vimentin staining (Figure 2B-D, Figure S3). In comparison to the control pericryptal mesenchyme cells, the mesenchymal cells carrying Lats1/2 deletion or YAP5SA expression were also highly proliferative, measured by Ki67 or PCNA staining (Figure 2E, F, Figure S3). Together, these data showed that YAP/TAZ activation drives over-proliferation of Gli1/PDGFR-expressing stromal cells and induces intestinal mesenchymal overgrowth.

**Figure 2.**
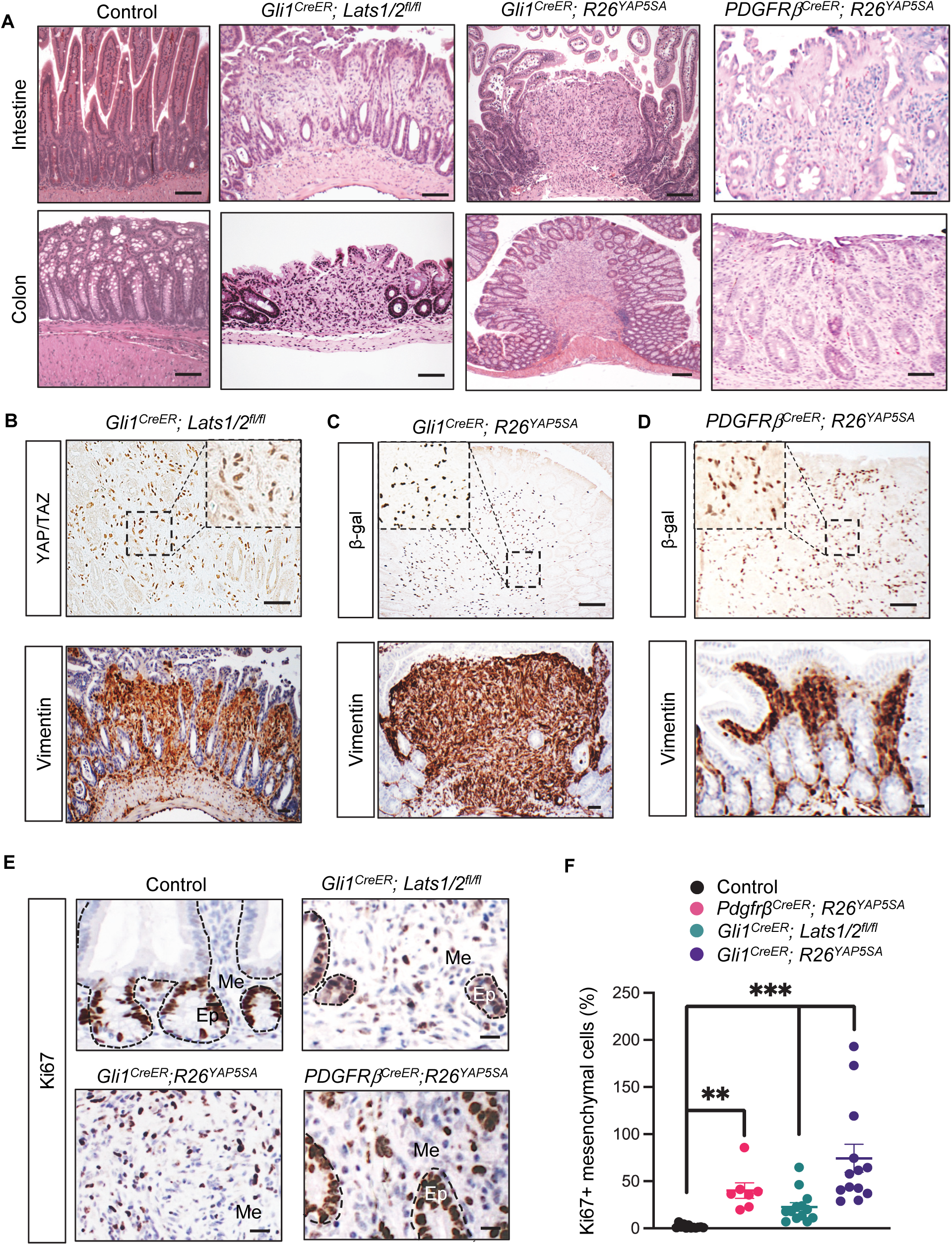
YAP/TAZ activation drives intestinal mesenchymal overgrowth. (**A**) Histology of intestine and colon of control, *Gli1*^*CreER*^; *Lats1/2*^*fl/fl*^, *Gli1*^*CreER*^; *R26*^*YAP5SA*^; *Pdgfrβ*^*CreER*^; *R26*^*YAP5SA*^. Scale bar, 50 μm. (**B**) Immunohistochemistry of YAP/TAZ and vimentin in mesenchymal overgrowths of *Gli1*^*CreER*^; *Lats1/2*^*fl/fl*^ mice. Inserts show staining in area indicated by dashed box. Scale bar 50 μm. (**C, D**) Immunohistochemistry of β-galactosidase (β-gal) and vimentin in mesenchymal overgrowths of *Gli1*^*CreER*^; *R26*^*YAP5SA*^ and *Pdgfrβ*^*CreER*^; *R26*^*YAP5SA*^ mice. Inserts show staining in area indicated by dashed box. Scale bar 50 μm. (**E**) Immunohistochemistry of Ki67 staining in control intestine and mesenchymal overgrowths in *Gli1*^*CreER*^; *Lats1/2*^*fl/fl*^, *Gli1*^*CreER*^; *R26*^*YAP5SA*^; *Pdgfrβ*^*CreER*^; *R26*^*YAP5SA*^ mice. Epithelia-mesenchyme boundaries are indicated by dashed lines. Me, mesenchyme; Ep, epithelia. Scale bar 50 μm. (**F**) Quantification of Ki67+ mesenchymal cells in control, *Pdgfrβ*^*CreER*^; *R26*^*YAP5SA*^, *Gli1*^*CreER*^; *Lats1/2*^*fl/fl*^, and *Gli1*^*CreER*^; *R26*^*YAP5SA*^. (n = 3 animals per group, 7-13 fields per group) Data are mean ± SEM. ** p<0.01, *** p<0.001. See also Figure S2, S3.

**Figure 3.**
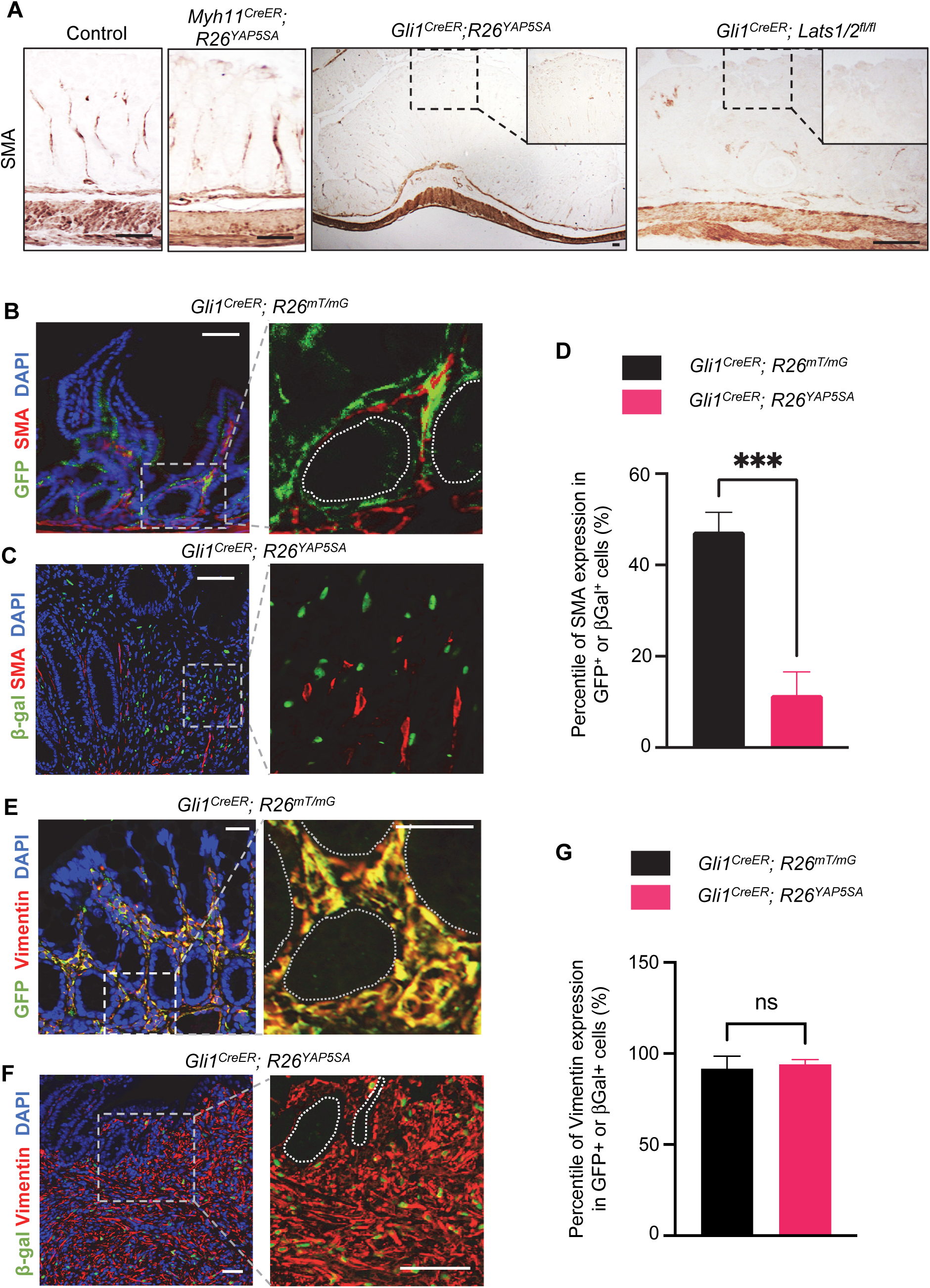
YAP/TAZ Activation Suppresses SMA Expression in Mesenchymal Progenitors. (**A**) Immunohistochemistry of SMA in the colon of control and *Myh11*^*CreER*^*;R26*^*YAP5SA*^ mice as well as overgrowths of *Gli1*^*CreER*^; *R26*^*YAP5SA*^ and *Gli1*^*CreER*^; *Lats1/2*^*fl/fl*^ mice. Scale bar, 50 μm. (**B**) Immunofluorescence staining of SMA and GFP in intestinal mesenchyme e of *Gli1*^*CreER*^; *R26*^*mT/mG*^ mice. Scale bar, 20 μm. (**C**) Immunofluorescence staining of SMA and β-galactosidase (β-gal) in mesenchymal overgrowth of *Gli1*^*CreER*^; *R26*^*YAP5SA*^ mice. (n = 6 polyps/areas per group) Scale bar, 20 μm. (**D**) Percentage of SMA expression in GFP+ or β-gal+ cells in control mesenchyme or overgrowths. Data are mean ± SEM. *** p<0.001. (**E**) Immunofluorescence staining of Vimentin and GFP in intestinal mesenchyme e of *Gli1*^*CreER*^; *R26*^*mT/mG*^ mice. Scale bar, 20 μm. (**F**) Immunofluorescence staining of Vimentin and β-galactosidase (β-gal) in mesenchymal overgrowth of *Gli1*^*CreER*^; *R26*^*YAP5SA*^ mice. Scale bar, 50 μm. Percentage of Vimentin expression in GFP+ or β-gal+ cells in control mesenchyme or overgrowths. (n = 3 polyps/areas per group) Data are mean ± SEM. ns, not significant.

### YAP/TAZ activation suppresses SMA expression in Gli1-expressing mesenchymal progenitors

Previous studies showed that the Gli1-expressing stromal cells located near the bases of intestinal crypts exhibit the characteristics of mesenchymal progenitor cells that are capable of undergoing differentiation of several lineages, including SMA-expressing myofibroblasts (Degirmenci et al., 2018). Interestingly, although it appears that YAP/TAZ activation did not affect SMA expression in the outer smooth muscle layers of *Myh11*^*CreER*^; *R26*^*YAP5SA*^ mice (Figure S1A), we noticed that SMA expression was markedly reduced in the mesenchymal overgrowths induced by *Gli1*^*CreER*^-induced YAP activation or Lats1/2 deletion, when compared to the stromal compartment in control animals (Figure 3A). These results suggested that YAP/TAZ activation may affect SMA expression in Gli1^+^ mesenchymal progenitors.

To further explore this phenotype, we generated the *Gli1*^*CreER*^; *R26*^*mT/mG*^ mice by crossing *Gli1*^*CreER*^ with the *R26*^*mT/mG*^ reporter allele (Muzumdar et al., 2007). *R26*^*mT/mG*^ is a cell membrane-targeted, two-color fluorescent Cre-reporter allele that expresses membrane-targeted Tomato (mT) prior to Cre-mediated excision and membrane-targeted GFP (mG) after excision (Muzumdar et al., 2007). Following TM injection in *Gli1*^*CreER*^; *R26*^*mT/mG*^ mice, we found that mGFP^+^ cells were readily detected in the pericryptal stromal regions (Figure 3B). Immunofluorescence staining showed that all GFP^+^ cells expressed Vimentin, a pan-mesenchymal marker, and only a subset of them were SMA positive (Figure 3B, D, E, G), consistent with the idea that these cells are likely the mesenchymal progenitors that can differentiate into SMA-expressing myofibroblasts (Degirmenci et al., 2018).

We calculated the percentage of Vimentin^+^ or SMA^+^ cells in GFP^+^ cells from the control G*li1*^*CreER*^; *R26*^*mT/mG*^ mice and compared it with the percentage of Vimentin^+^ or SMA^+^ cells in β-galactosidase^+^ YAP5SA-expressing mutant cells in the mesenchymal overgrowth from *Gli1*^*CreER*^; *R26*^*YAP5SA*^ mice (Figure 3B-G). As expected, almost all control and YAP5SA-expressing cells expressed Vimentin (Figure 3E-G), whereas immunofluorescence staining of β-galactosidase and SMA in mesenchymal overgrowths showed very little co-expression in cells, suggesting that YAP activation in the Gli1^+^ cells may potentially inhibit SMA expression (Figure 3B-D). Together, these results indicated that YAP activation does not influence SMA expression in the terminally differentiated smooth muscle; however, it may alter the differentiation of the Gli1^+^ mesenchymal progenitors by suppressing SMA expression.

### Mesenchymal YAP/TAZ activation promotes Wnt ligand production in the stem cell niche

It has been demonstrated that Gli1-expressing stromal cells in the stem cell niche serve as one of the major sources of Wnt ligands and agonists and are essential for maintaining the epithelial stem cells (Degirmenci et al., 2018; Valenta et al., 2016). We set out to examine whether Hippo-YAP signaling regulates Wnt expression in the Gli1^+^ cells. First, we measured the mRNA levels of Wnts in isolated Gli1^+^ mesenchymal cells with and without YAP5SA expression. We found that YAP5SA expression induced significantly higher levels of several Wnt ligands and agonists, including *Wnt2, Wnt2b, Wnt4, Rspo1*, and *Rspo3* (Figure 4A), suggesting an important role of Hippo-YAP signaling in controlling Wnt production in the stem cell niche.

**Figure 4.**
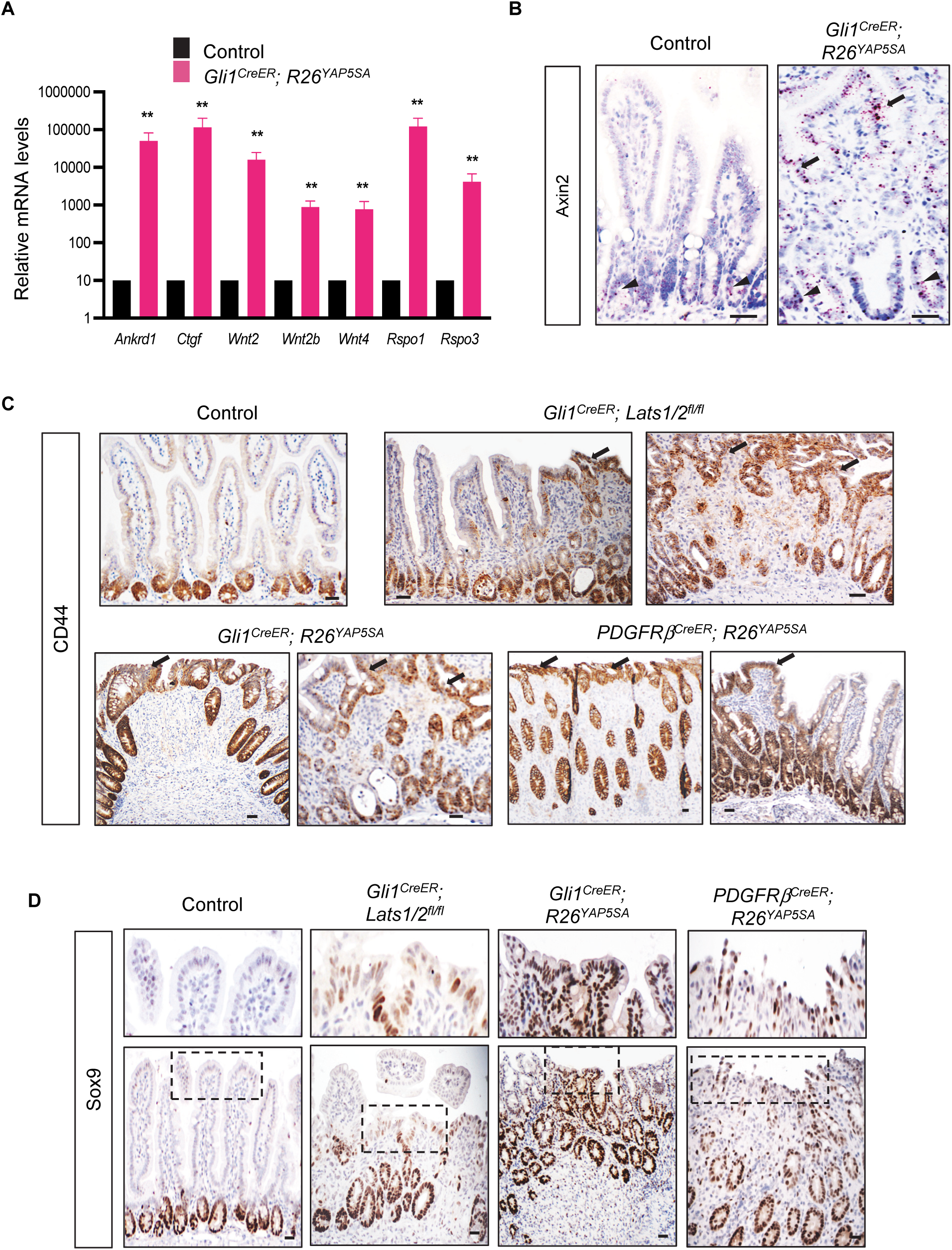
Mesenchymal YAP/TAZ Activation Promotes Epithelial Wnt Pathway Activation. **(A)**Real-time PCR analysis of YAP targets *Ankrd1* and *Cyr61* and Wnt ligands/agonists *Wnt2b, Wnt4, Rspo1, and Rspo3* mRNA levels in control and *Gli1*^*CreER*^*;R26*^*YAP5SA*^ mesenchyme. (technical triplicate per sample, n= 5 animals per group) Data are mean ± SEM., ** p<0.01. (**B**) RNAscope analysis of *Axin2* mRNA expression in the epithelia of control and *Gli1*^*CreER*^*;R26*^*YAP5SA*^ mice. Arrowheads point to Axin2 expression in the crypt, and arrows point to Axin2 expression in the epithelia adjacent to mesenchymal overgrowths. Scale bar, 20 μm. (**C**) Immunohistochemistry of CD44 in control, *Gli1*^*CreER*^*;Lats1/2*^*fl/fl*^, *Gli1*^*CreER*^*;R26*^*YAP5SA*^; and *Pdgfrβ*^*CreER*^*;R26*^*YAP5SA*^ epithelia. Arrows point to CD44 expression in the epithelia adjacent to mesenchymal overgrowths. Scale bar, 20 μm. (**D**) Immunohistochemistry of SOX9 in control, *Gli1*^*CreER*^*;Lats1/2*^*fl/fl*^, *Gli1*^*CreER*^*;R26*^*YAP5SA*^; and *Pdgfrβ*^*CreER*^*;R26*^*YAP5SA*^ epithelia. Upper panels show areas indicated by dashed boxes. Scale bar, 20 μm.

To further explore the effect of mesenchymal YAP/TAZ activation on epithelial Wnt signaling activity, we measured the expression levels of well-established intestinal Wnt target genes, including *Axin2, Cd44* and *Sox9*, in the overgrowths detected in *Gli1*^*CreER*^; *Lats1/2*^*fl/fl*^, *Gli1*^*CreER*^;*R26*^*YAP5SA*^ and *Pdgfrb*^*CreER*^*;R26*^*YAP5SA*^ mice following Tamoxifen injection. RNAscope for Axin2 and IHC for both CD44 and Sox9 showed that all these Wnt targets were highly expressed in the epithelia immediately adjacent to areas of mesenchymal overgrowths, in addition to the crypt where Wnt signaling is normally activated (Figure 4B-D). Altogether, these data suggested that YAP/TAZ activation in Gli1^+^ mesenchymal cells promotes Wnt pathway activation in the adjacent epithelia, thereby regulating mesenchymal-epithelial crosstalk in the intestine.

### Mesenchymal YAP activation promotes intestinal epithelial regeneration induced by DSS treatment

Wnt signaling is crucial for stem cell renewal and regeneration following injury (Kretzschmar and Clevers, 2017; Ma et al., 2019; Mah et al., 2016; Nusse and Clevers, 2017). Since our data suggested that YAP activation within the mesenchymal compartment upregulates epithelial Wnt signaling (Figure 4), we hypothesized that mesenchymal YAP activation may expediate intestinal regeneration after injury.

To test this hypothesis, we utilized the intestinal injury model induced by the compound dextran sodium sulfate (DSS), regularly used *in vivo* to model colitis in mice (Chassaing et al., 2014). DSS-induced injury is characterized by massive epithelial death, impairment of glandular architecture, and inflammation. We treated adult male animals with 2.5% DSS in their drinking water for 7 days, after which we’d replace DSS water with regular water and allow an additional 14 days for recovery (Figure 5A). Pathological evaluation revealed severe epithelial damage following 7 days of DSS treatment (Day 7), in which the wounded areas exhibited an absence of glandular architecture and no epithelial layer (Figure 5C). As recovery progresses, colonic mucosa exhibited would healing activities including the formation of nascent crypts flanking the wound, and epithelial differentiation at the surface of the wound (Figure 5C). IHC staining of CD44 showed that the nascent crypts exhibited high Wnt pathway activation, and terminal differentiation of epithelial cells during recovery was revealed by the staining of Keratin 20 (Figure 5B). Interestingly, we noticed that mesenchymal expression of YAP/TAZ during injury and subsequent recovery was also significantly higher than its levels in the mesenchyme before injury or after recovery (Figure 5B), suggesting an intriguing possibility that mesenchymal YAP/TAZ activity may potentially aid in epithelial regeneration.

**Figure 5.**
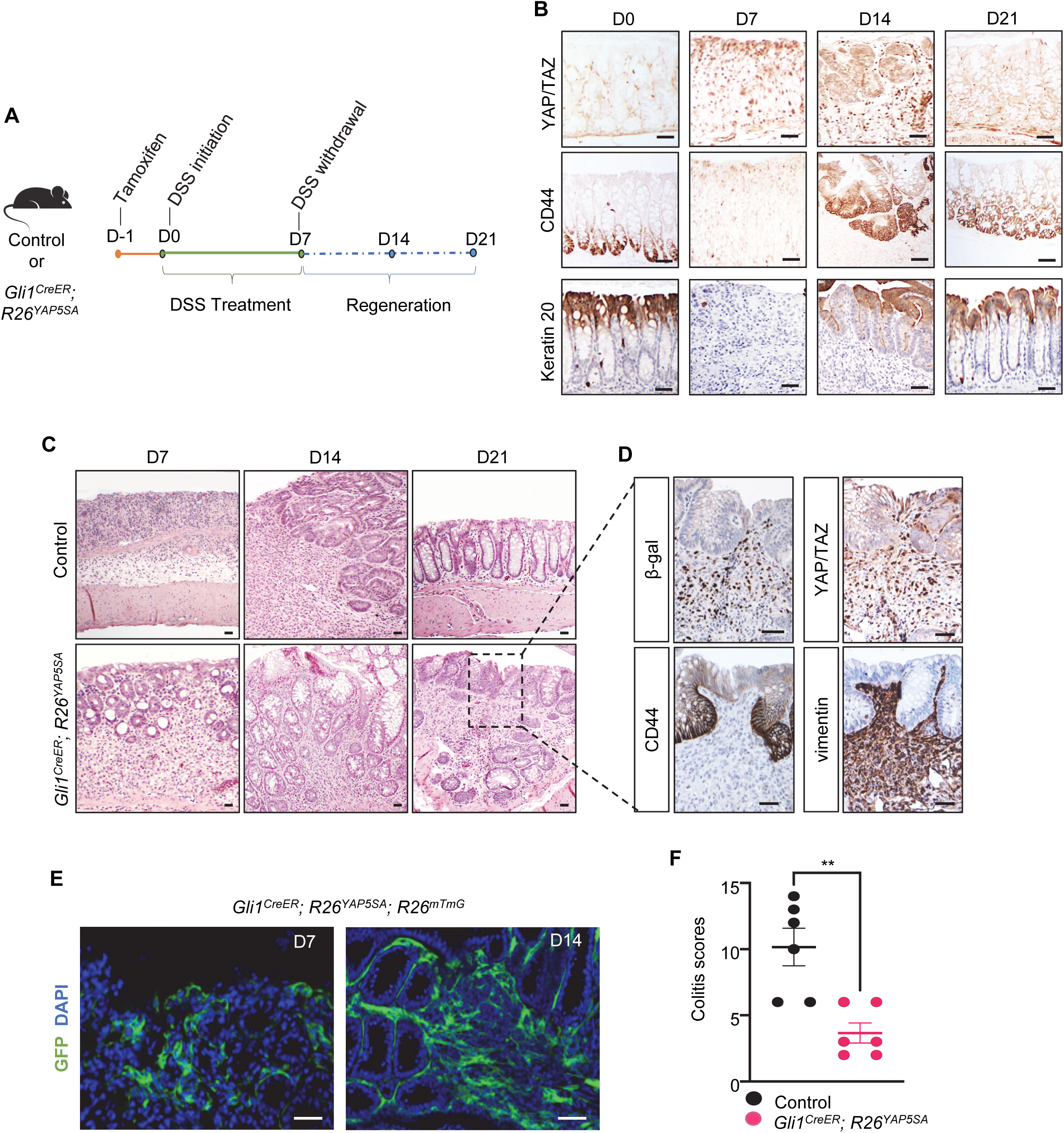
Mesenchymal YAP activation promotes intestinal regeneration induced by DSS treatment. **(A)**Schematic of experimental design, indicating tamoxifen treatment, duration of DSS treatment and recovery. (**B**) Immunohistochemistry of YAP/TAZ, CD44, and Keratin-20 in control intestine during DSS treatment and subsequent recovery at timepoints Day 0 (D0), Day 7 (D7), Day 14 (D14) and Day 21 (D21). Scale bar, 20 μm. (**C**) Histology of control and *Gli1*^*CreER*^; *R26*^*YAP5SA*^ colon at timepoints D7, D14, and D21 of recovery. Scale bar, 20 μm. (**D**) Immunohistochemistry of β-galactosidase (β-gal), YAP/TAZ, CD44, and vimentin in areas of mesenchymal overgrowth in *Gli1*^*CreER*^; *R26*^*YAP5SA*^ colon at D21. Scale bar, 20 μm. (**E**) GFP signal in *Gli1*^*CreER*^; *R26*^*YAP5SA*^; *R26*^*mT/mG*^ colon at D7 and D14 during regeneration. Scale bar, 50 μm. (**F**) Colitis scoring of control and *Gli1*^*CreER*^; *R26*^*YAP5SA*^ colon. (n=4 animals per group). Data are mean ± SEM., ** p<0.01.

Prior report has also demonstrated that Wnt-secreting Gli1^+^ mesenchymal cells not only serve as the essential niche for the intestinal stem cells at the normal state, but they are also markedly expanded after DSS treatment, indicating their involvement in epithelial recovery from injury (Degirmenci et al., 2018). Thus, to examine the effect of YAP activation in the Gli1^+^ mesenchymal cells on intestinal regeneration, *Gli1*^*CreER*^; *R26*^*YAP5SA*^ or *Gli1*^*CreER*^; *R26*^*YAP5SA*^; *R26*^*mT/mG*^ mice were injected with Tamoxifen to induce Cre recombination followed by DSS treatment. We found that, compared to control mice, *Gli1*^*CreER*^; *R26*^*YAP5SA*^ mice showed significantly more robust recovery from injury. At Day 7, nascent crypts and epithelial layers were absent in the wounded area in control mice but readily detected in *Gli1*^*CreER*^; *R26*^*YAP5SA*^ mice (Figure 5B). During the recovery, epithelial regeneration was also more advanced in *Gli1*^*CreER*^; *R26*^*YAP5SA*^ mice (Figure 5B). At Day 21 when colon homeostasis was restored in control mice, we detected mesenchymal overgrowths in *Gli1*^*CreER*^; *R26*^*YAP5SA*^ mice, probably due to the persistent over-expression of YAP5SA in the Gli1^+^ cells after recovery from injury (Figure 5B). Consistent with the previous report of the increase of Gli1^+^ cells in DSS-treated mice (Degirmenci et al., 2018), we also found that YAP5SA-expressing/GFP+ mesenchymal cells in *Gli1*^*CreER*^; *R26*^*YAP5SA*^; *R26*^*mT/mG*^ mice were present at the wounded areas, adjacent to the epithelia surrounding nascent colon crypts that expressed the Wnt target gene CD44 (Figure 5D). In addition, we histologically scored the severity of DSS-mediated injury (also referred to as colitis) in which inflammation, regeneration, crypt damage, and percent involvement were assessed in the colon for both control and mutant groups using a previously published grading system (Dieleman et al., 1998), in which, an individual total score ranges from 0-14 (least severe to most severe). Our analysis revealed that *Gli1*^*CreER*^; *R26*^*YAP5SA*^ mutant colons were significantly less injured than control immediately following DSS treatment and during recovery (Figure 5E). Altogether, our data revealed mesenchymal YAP/TAZ upregulation during DSS-induced injury, and we showed that YAP activation in the niche-forming Gli1^+^ mesenchymal cells promotes intestinal epithelial regeneration after injury.

## DISCUSSION

The Hippo-YAP signaling pathway is intimately associated with intestinal homeostasis and tumorigenesis (Gregorieff and Wrana, 2017; Hong et al., 2016; Zheng and Pan, 2019); however, its precise function remains elusive. Much of the focus has been on Hippo-YAP signaling in intestinal epithelium, and little is known about its function in the mesenchyme, a tissue layer that plays an essential role during intestinal development and homeostasis. Our recent report linked Hippo signaling to the embryonic gut mesenchyme, where YAP activation drives the expansion of the mesenchymal progenitor populations (Cotton et al., 2017). Our study here focused on the role of Hippo-Yap signaling in adult intestinal mesenchymal cells. Recent studies have demonstrated that several populations of subepithelial mesenchymal cells, including the Gli1^+^ progenitor cells that form the essential Wnt-secreting niche to sustain intestinal stem cells (Degirmenci et al., 2018; Greicius et al., 2018; Shoshkes-Carmel et al., 2018). However, how these cells are regulated is largely unexplored. We identified Hippo/YAP as a key signaling pathway that controls proliferation and differentiation of these niche-forming mesenchymal cells and regulates Wnt ligand production during intestinal epithelial-mesenchymal crosstalk.

We found that Lats1/2 deletion or YAP activation in Gli1^+^ cells induced drastic mesenchymal overgrowth, which is in contrast to the effect of Hippo pathway disruption in the smooth muscle layer, where YAP/TAZ activation has no apparent growth effect (Figure 1). It is probably in part due to the baseline YAP/TAZ activity in different mesenchymal populations where YAP/TAZ expression is at a much higher level in the smooth muscle layer than in the subepithelial stromal compartment (Figure 1). It is also intriguing that YAP/TAZ activation has a differential effect on SMA expression in these different mesenchymal populations. Our result showed that YAP activation suppresses SMA expression in Gli1^+^ progenitors. It is consistent with our prior report that YAP activation prevents SMA induction in the primitive embryonic mesenchyme (Cotton et al., 2017). However, neither Lats1/2 deletion or YAP overaction can suppress SMA expression in the outer smooth muscle layer. Thus, it is interesting to speculate that different wiring of the genetic or epigenetic circuitries in terminally differentiated smooth muscle cells and mesenchymal progenitors may alter the ability of YAP/TAZ to modulate the expression of certain sets of genes related to cell differentiation. Clearly, further investigation is needed to address these issues.

The interaction between Wnt and Hippo-YAP signaling in the intestine has been investigated; however, most of the reports addressed the crosstalk of signal transduction of these two pathways in the epithelial stem cells (Cai et al., 2015; Cheung et al., 2020; Gregorieff and Wrana, 2017; Hong et al., 2016; Li et al., 2020; Zanconato et al., 2016). Our study here identified a role of Hippo-YAP signaling on Wnt ligand production in pericryptal mesenchymal cells, uncovering a critical molecular mechanism on how the precise levels of Wnt ligands are controlled within the stem cell niche. Thus, in addition to demonstrating the mesenchymal Hippo pathway as an essential niche signal, our work identified another layer of the complex Hippo-Wnt functional interactions that involve both the epithelial and mesenchymal compartments during intestinal homeostasis.

Our results revealed that mesenchymal Hippo-YAP signaling not only regulates normal homeostasis but also contributes to intestinal pathogenesis. We found that the YAP/TAZ expression level in the mesenchyme is markedly up-regulated during DSS-induced intestinal injury and subsequent recovery and demonstrated that mesenchymal YAP activation promotes intestinal regeneration in a mouse model of colitis. These data are consistent with our result that mesenchymal Hippo-YAP controls Wnt production in the stem cell niche, and it also links mesenchymal Hippo-YAP regulation to intestinal pathogenesis. Hippo signaling is known to interact with other signaling pathways, including receptor tyrosine kinases and TGF/BMP that have been shown to function in the stem cell niche during intestinal hemostasis (McCarthy et al., 2020). It is interesting to explore how the interplays between Hippo-YAP and these signaling pathways in the mesenchyme may coordinate intestinal epithelial-mesenchymal crosstalk and how they can be exploited for the design of new therapeutic strategy towards treatment of intestinal diseases and cancers.

## MATERIALS AND METHODS

### Key resource table

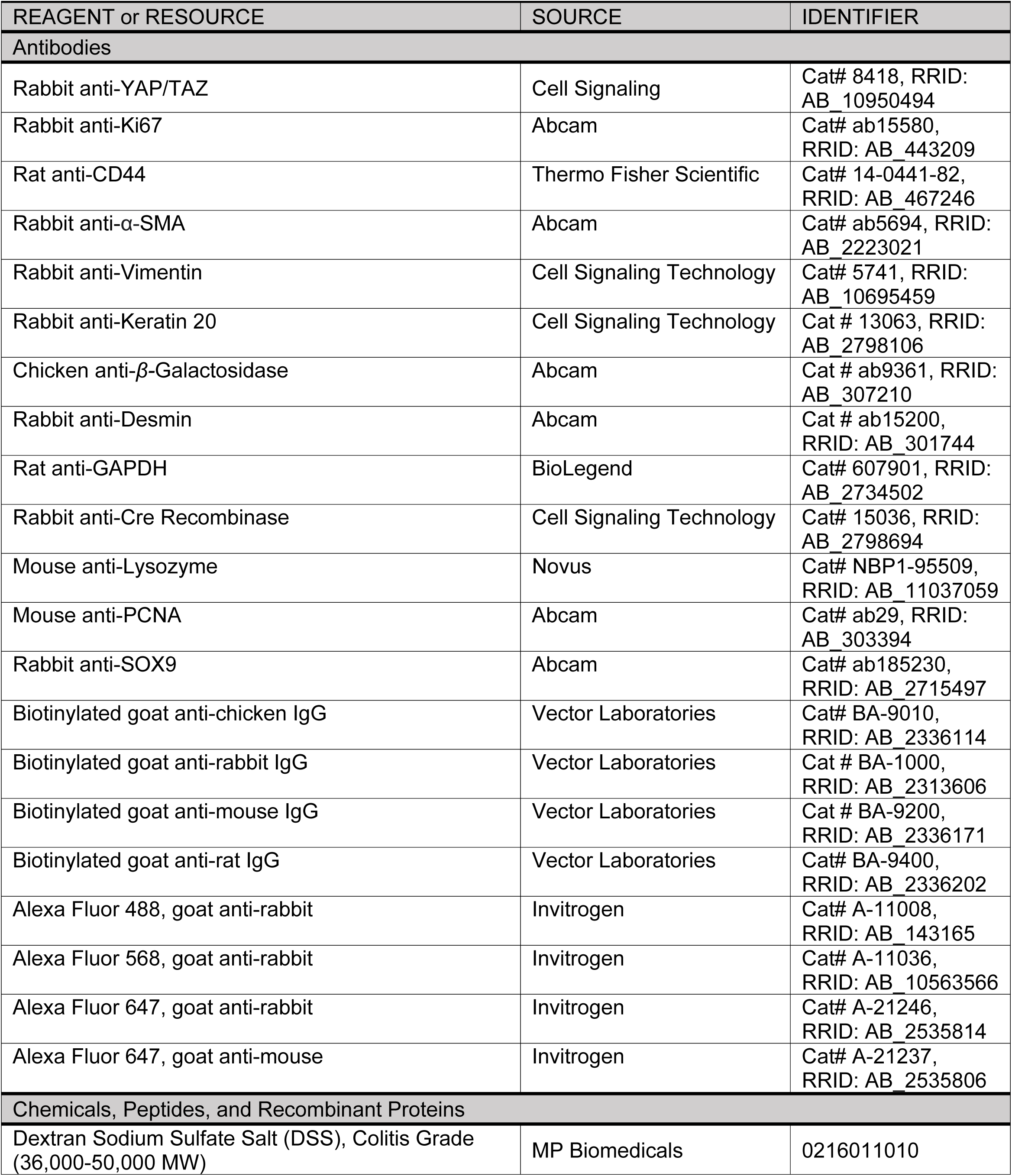
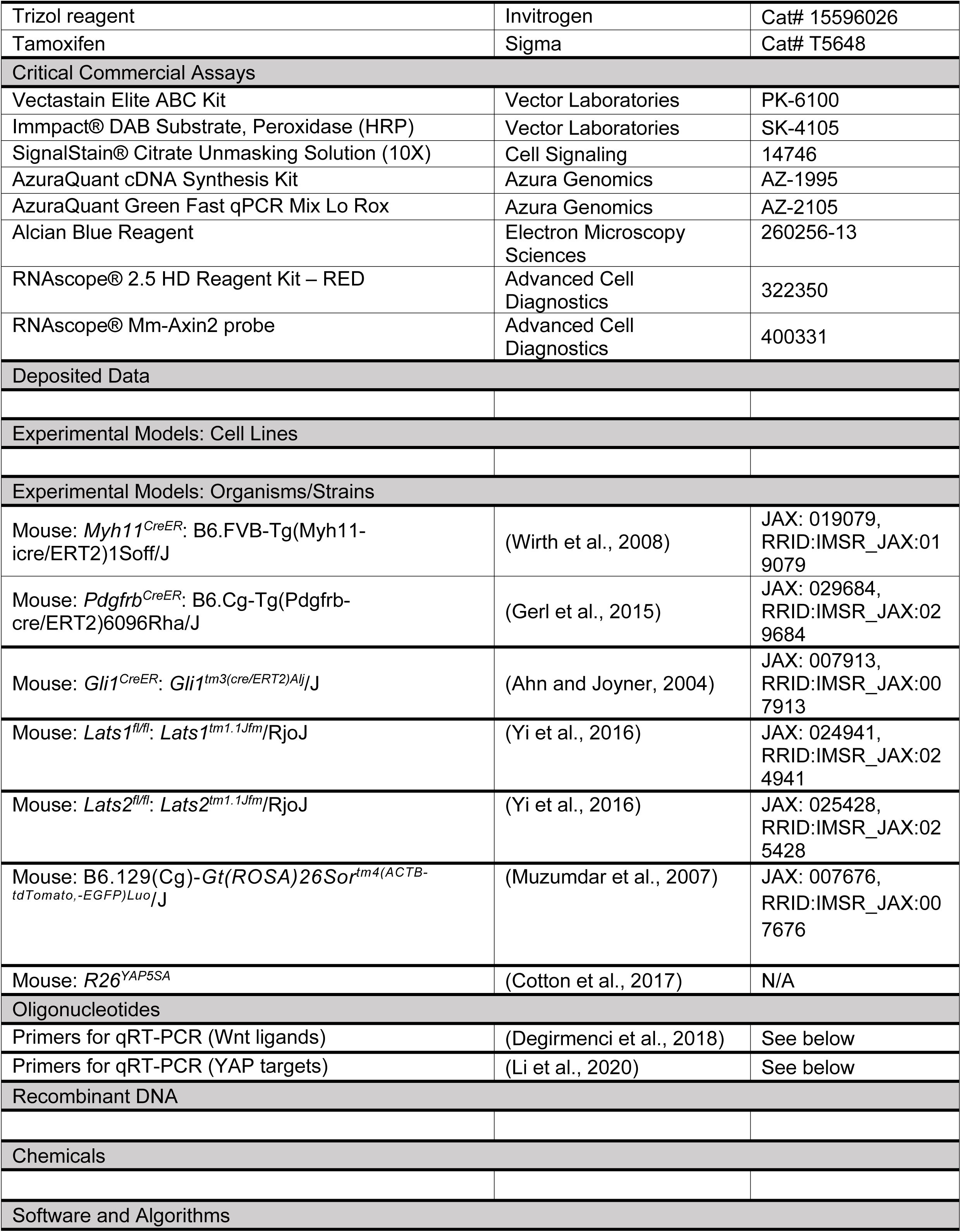
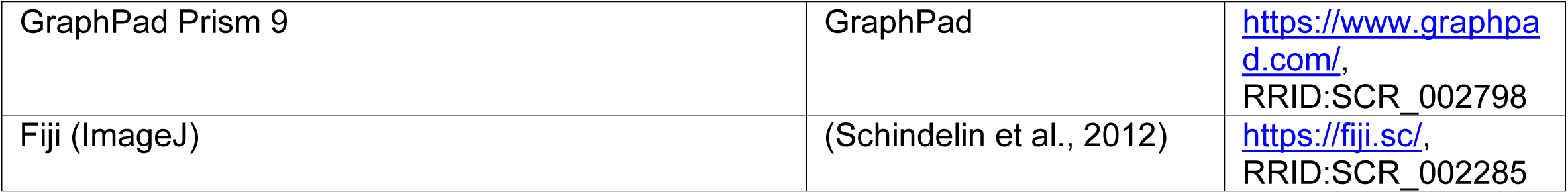

#### Mice

All animals use protocols were reviewed and approved by The University of Massachusetts Medical School Institutional Animal Care and Use Committee. *Gli1*^*CreER*^ (Ahn and Joyner, 2004), *Myh11*^*CreER*^ (Wirth et al., 2008), *PdgfrβCreER* (Gerl et al., 2015), *Lats1*^*flox*^ and *Lats2*^*flox*^ (Yi et al., 2016), and *R26*^*mT/mG*^ (Muzumdar et al., 2007) mice were obtained from the Jackson laboratory. *R26*^*YAP5SA*^ mice were described previously (Cotton et al., 2017). Cre activation of the inducible Cre lines was achieved by one-time intraperitoneal injection of 120mg/kg Tamoxifen (Sigma) into one-month old mice with appropriate genotypes.

#### DSS-colitis model

*Gli1*^*CreERA*^; *R26*^*mT/mG*^ control and *Gli1*^*CreER*^; *R26*^*YAP5SA*^; *R26*^*mT/mG*^ mutant animals of 8 weeks of age or older were induced with one single dose of 300 mg/kg tamoxifen one day prior to treatment with 2.5% w/v dextran sodium sulfate (DSS) in autoclaved drinking water. Animals were treated for 7 consecutive days before regular drinking water was re-introduced. Animals from each group were sacrificed at pre-determined time points: Immediately following DSS treatment (D7), 7 days following DSS treatment (D14), and 14 days following DSS treatment (D21). The colon was harvested and prepared for both paraffin and OCT embedding and stained with hematoxylin and eosin as described under “Tissue Collection and Histology.” Colitis injury was histologically scored using H&E paraffin sections and criteria previously published (Dieleman et al., 1998).

#### Tissue Collection and Histology

Following euthanasia, mouse intestinal or polyp tissue was dissected and fixed in 10% Neutral Buffered Formalin (NBF) at 4°C overnight. For paraffin sections, tissue was dehydrated, embedded in paraffin, and sectioned at 6 µm. For frozen sections, tissue was dehydrated in 30% sucrose overnight at 4°C, embedded in OCT, and sectioned at 12 µm. Paraffin sections were stained using standard hematoxylin & eosin reagents. For intestinal epithelium and mesenchyme isolation, mouse small intestinal tissues at different postnatal stages are dissected and washed in cold PBS, before transferring to PBS containing 3mM EDTA for rotation at 4°C for 30 mins. After vigorous shaking for 2 mins, the epithelial tissues are collected in the supernatant, while the remaining mesenchymal tissues are washed and incubated with the digestion buffer containing Collagenase XI and Dispase at 37°C for 30min, and the samples are then subjected to western blot analysis.

#### Immunohistochemistry and Immunofluorescence

For immunohistochemistry (IHC), sections were deparaffinized and rehydrated before undergoing heat-induced antigen retrieval in 10mM sodium citrate buffer (pH 6.0) for 30 minutes. Slides were blocked for endogenous peroxidase for 20 minutes, then blocked for 1 hour in 5% BSA, 1% goat serum, 0.1% Tween-20 buffer in PBS, and incubated overnight at 4°C in primary antibody diluted in blocking buffer or SignalStain® Antibody Diluent (Cell Signaling). Slides were incubated in biotinylated secondary antibodies for 1 hour at room temperature and signal was detected using the Vectastain Elite ABC kit (Vector Laboratories). For immunofluorescence (IF), cells or tissue sections were fixed by 4% paraformaldehyde for 5 minutes, blocked for 1 hour and incubated overnight at 4°C in primary antibody diluted in blocking buffer. Slides were then incubated for 1 hour at room temperature in Alexa Fluor-conjugated secondary antibodies (Invitrogen) at 1:500 dilution in blocking buffer and mounted using mounting media with DAPI (EMS).

Primary antibodies used for IHC/IF were YAP/TAZ (Cell Signaling), Vimentin (Cell Signaling), Ki67 (Abcam), CD44 (eBioscience), β-galactosidase (Abcam), Desmin (Abcam), SOX9 (Abcam), Keratin 20 (Cell Signaling), lysozyme (Novus), PCNA (Abcam), Cre recombinase (Cell signaling), and α-smooth muscle actin (SMA) (Abcam).

#### Western blot analysis

Freshly isolated mouse tissue was lysed in lysis buffer (50 mM Tris-HCl, pH 7.4, 150 mM NaCl, 0.5 mM EDTA, 1% Triton X-100, phosphatase inhibitor cocktail, cOmplete EDTA-free protease inhibitors cocktail) for 30min at 4°C. The supernatants of the extracts were then used for western blot following the protocols described previously (Cotton et al., 2017) and the primary antibodies used in these assays were listed as follows: YAP/TAZ (Cell Signaling) and GAPDH (Bethyl). HRP-conjugated Secondary antibodies were obtained from Jackson Laboratories.

#### Quantitative Real-Time PCR

Total RNA was isolated using Trizol reagent (Invitrogen). cDNA synthesis was performed using AzuraQuant™ cDNA synthesis kit (Azura Genomics), and the number of transcripts were quantified using AzuraQuant™ Green Fast qPCR Mix Lo-Rox (Azura Genomics), with the respective oligonucleotides: (CTGF forward: TGTGCACTGCCAAAGATGGTGCAC, reverse: TGGGCAGGCGCACGTCCATG; Cyr61 forward: GAGGCTTCCTGTCTTTGGCAC, reverse: ACTCTGGGTTGTCATTGGTAAC; ANKRD1 forward: GGAACAACGGAAAAGCGAGAA, reverse: GAAACCTCGGCACATCCACA) in QuantStudio 6 Flex Real-Time PCR Systems (Applied Biosystems). All qPCR experiments were conducted in biological triplicates, error bars represent mean ± standard error mean, and Student’s t-test was used to generate p-values (* = p value <0.05; ** = p value <0.01).

#### Statistical analysis

No statistical method was used to predetermine sample size. The experiments were not randomized. For biochemical experiments we performed the experiments at least three independent times. Experiments for which we showed representative images were performed successfully at least 3 independent times. No samples or animal were excluded from the analysis. The investigators were not blinded to allocation during experiments and outcome assessment. Student’s t-test was used to generate p-values (* = p value ≤0.05; ** = p value ≤0.01). The variance was similar between groups that we compared

## ACKNOWLEDGEMENT

This work was supported by grants from National Institutes of Health R01DK127180 and R01DK127207 to J.M. and R01CA238270 and R01DK107651 to X.W, and in part by Samuel M. Fisher Memorial–MRA (Melanoma Research Alliance) Established Investigator Award to X.W. We also thank the members of Wu lab and Mao lab for helpful discussion and comments on the manuscript.

## AUTHOR CONTRIBUTIONS

KD and JM conceived and designed the study. KD, AS, JLC, ZT, HL and LJZ acquired the data. KD, XW and JM analyzed and interpreted the data. KD and JM wrote the manuscript. JM supervised the study.

### COMPETING INTERESTS

The authors declare no competing interest.

---

**Figure S1.**
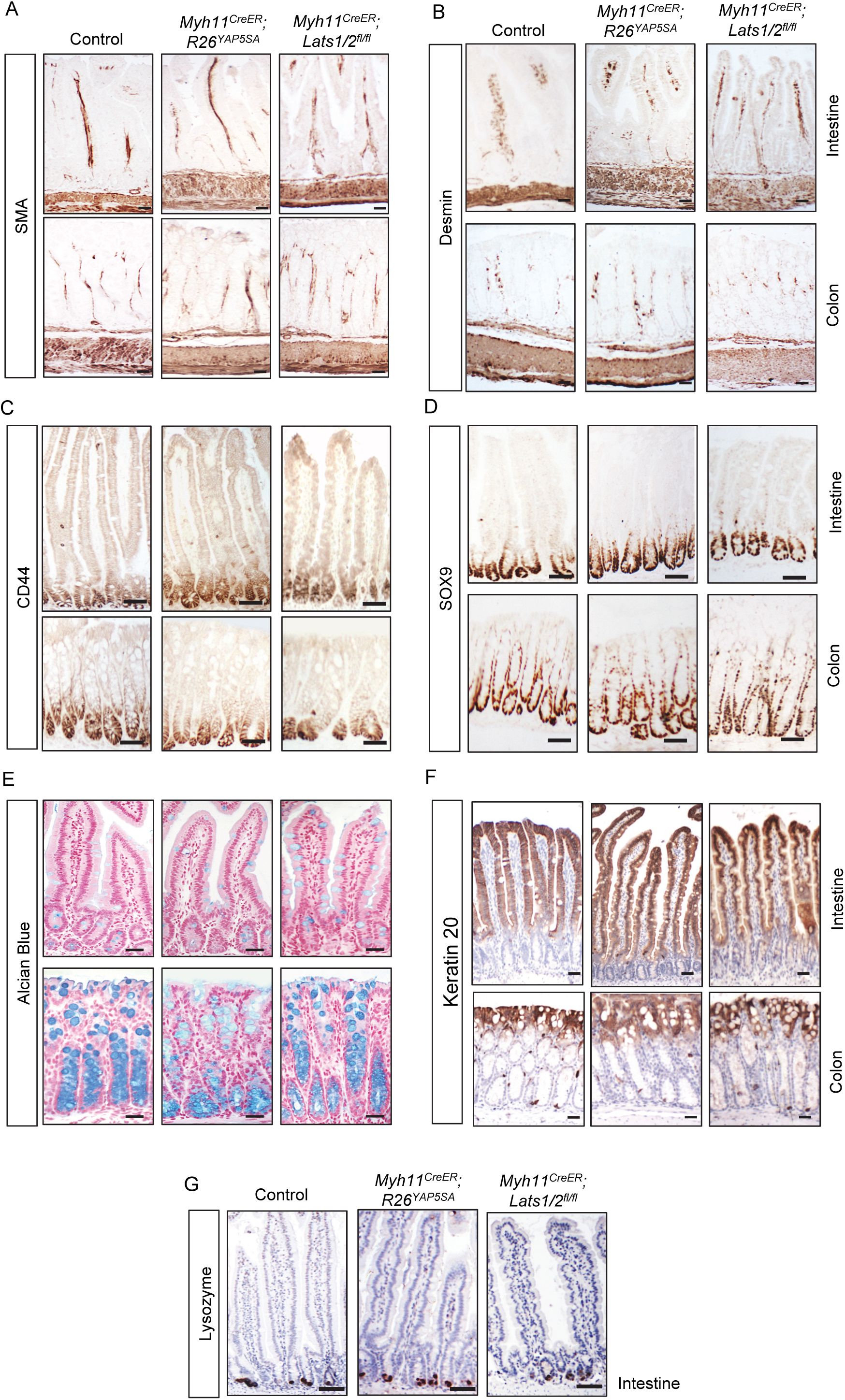
Intestinal growth and differentiation in *Myh11*^*CreER*^; *R26*^*YAP5SA*^ and *Myh11*^*CreER*^; *Lats1/2*^*fl/fl*^ mice. (**A**-**G**) Immunohistochemistry of SMA, Desmin, CD44, Sox9, Keratin 20 and Lysozyme, and Alcian blue staining in intestine and colon of control, *Myh11*^*CreER*^; *R26*^*YAP5SA*^, and *Myh11*^*CreER*^; *Lats1/2*^*fl/fl*^ animals. Scale bar, 20 μm.

**Figure S2.**
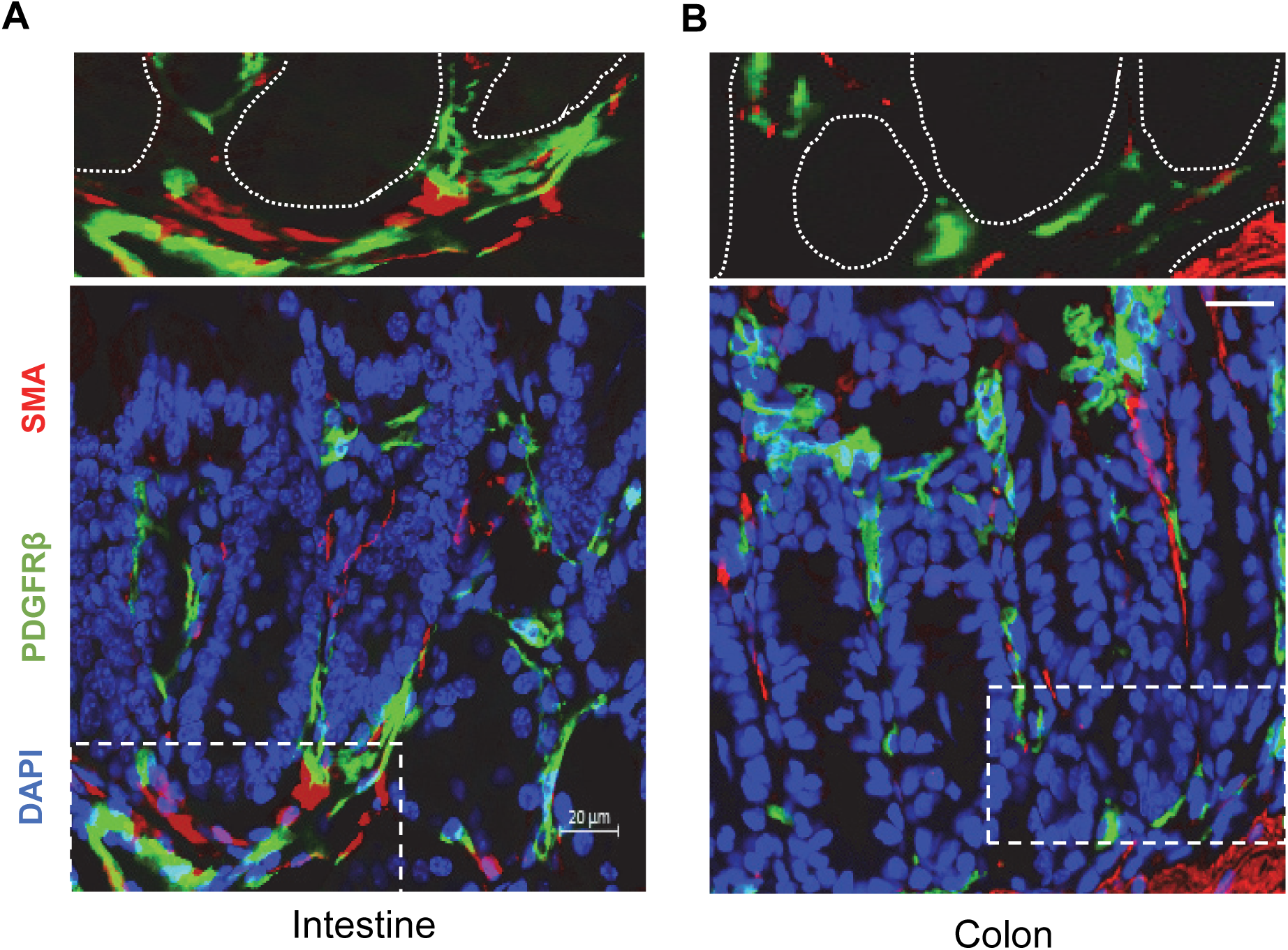
PDGFRβ expression in subepithelial mesenchyme adjacent to the crypt. (**A, B**) Immunofluorescence of Pdgfrβ and SMA in control intestine and colon. Top panels show Pdgfrβ and SMA staining indicated by dashed box. Scale bar, 20 μm.

**Figure S3.**
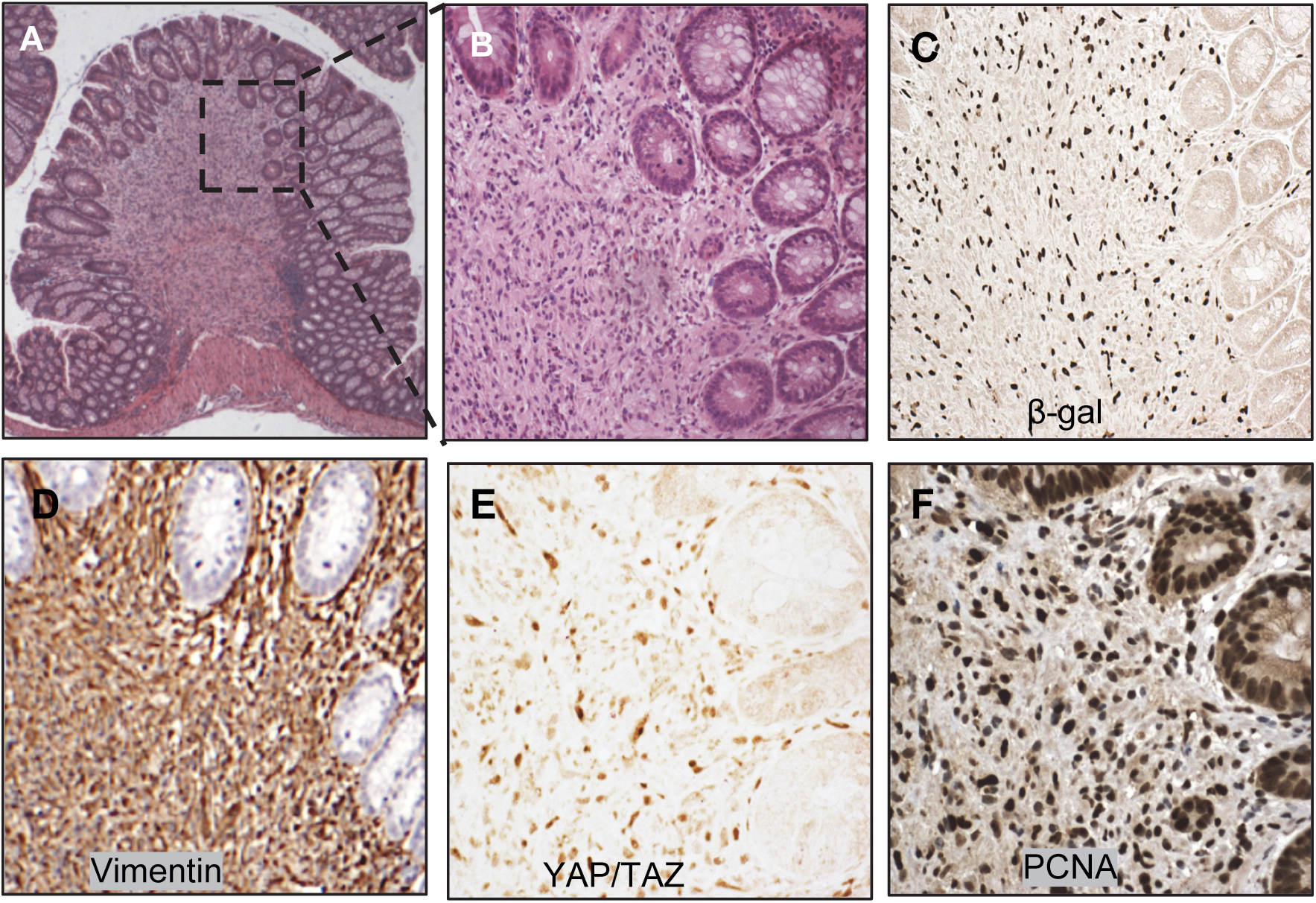
YAP/TAZ activation drives colon mesenchymal overgrowth. (**A**) Histology of mesenchymal overgrowth in *Gli1*^*CreER*^; *R26*^*YAP5SA*^ colon. Scale bar, 50 μm. (**B-F**) Histology and immunohistochemistry of β-galactosidase (β-gal), Vimentin, YAP/TAZ and PCNA of area indicated by dashed box in (A).

## Notes

### Competing Interest Statement

The authors have declared no competing interest.

## REFERENCES

Ahn, S., and Joyner, A.L. (2004). Dynamic changes in the response of cells to positive hedgehog signaling during mouse limb patterning. Cell 118, 505–516.

Ahn, S., and Joyner, A.L. (2005). In vivo analysis of quiescent adult neural stem cells responding to Sonic hedgehog. Nature 437, 894–897.

Barry, E.R., Morikawa, T., Butler, B.L., Shrestha, K., de la Rosa, R., Yan, K.S., Fuchs, C.S., Magness, S.T., Smits, R., Ogino, S., et al. (2013). Restriction of intestinal stem cell expansion and the regenerative response by YAP. Nature 493, 106–110.

Cai, J., Maitra, A., Anders, R.A., Taketo, M.M., and Pan, D. (2015). beta-Catenin destruction complex-independent regulation of Hippo-YAP signaling by APC in intestinal tumorigenesis. Genes Dev 29, 1493–1506.

Cai, J., Zhang, N., Zheng, Y., de Wilde, R.F., Maitra, A., and Pan, D. (2010). The Hippo signaling pathway restricts the oncogenic potential of an intestinal regeneration program. Genes Dev 24, 2383–2388.

Chassaing, B., Aitken, J.D., Malleshappa, M., and Vijay-Kumar, M. (2014). Dextran sulfate sodium (DSS)-induced colitis in mice. Curr Protoc Immunol 104, 15.25.11-15.25.14.

Cheung, P., Xiol, J., Dill, M.T., Yuan, W.C., Panero, R., Roper, J., Osorio, F.G., Maglic, D., Li, Q., Gurung, B., et al. (2020). Regenerative Reprogramming of the Intestinal Stem Cell State via Hippo Signaling Suppresses Metastatic Colorectal Cancer. Cell Stem Cell 27, 590–604 e599.

Cotton, J.L., Li, Q., Ma, L., Park, J.S., Wang, J., Ou, J., Zhu, L.J., Ip, Y.T., Johnson, R.L., and Mao, J. (2017). YAP/TAZ and Hedgehog Coordinate Growth and Patterning in Gastrointestinal Mesenchyme. Dev Cell 43, 35–47 e34.

Degirmenci, B., Valenta, T., Dimitrieva, S., Hausmann, G., and Basler, K. (2018). GLI1-expressing mesenchymal cells form the essential Wnt-secreting niche for colon stem cells. Nature 558, 449–453.

Dieleman, L.A., Palmen, M.J., Akol, H., Bloemena, E., Peña, A.S., Meuwissen, S.G., and Van Rees, E.P. (1998). Chronic experimental colitis induced by dextran sulphate sodium (DSS) is characterized by Th1 and Th2 cytokines. Clin Exp Immunol 114, 385–391.

Gerl, K., Miquerol, L., Todorov, V.T., Hugo, C.P., Adams, R.H., Kurtz, A., and Kurt, B. (2015). Inducible glomerular erythropoietin production in the adult kidney. Kidney Int 88, 1345–1355.

Gregorieff, A., Liu, Y., Inanlou, M.R., Khomchuk, Y., and Wrana, J.L. (2015). Yap-dependent reprogramming of Lgr5(+) stem cells drives intestinal regeneration and cancer. Nature 526, 715–718.

Gregorieff, A., and Wrana, J.L. (2017). Hippo signalling in intestinal regeneration and cancer. Curr Opin Cell Biol 48, 17–25.

Greicius, G., Kabiri, Z., Sigmundsson, K., Liang, C., Bunte, R., Singh, M.K., and Virshup, D.M. (2018). PDGFRalpha(+) pericryptal stromal cells are the critical source of Wnts and RSPO3 for murine intestinal stem cells in vivo. Proc Natl Acad Sci U S A 115, E3173–E3181.

Halder, G., and Camargo, F.D. (2013). The hippo tumor suppressor network: from organ size control to stem cells and cancer. Cancer Res 73, 6389–6392.

Hong, A.W., Meng, Z., and Guan, K.L. (2016). The Hippo pathway in intestinal regeneration and disease. Nat Rev Gastroenterol Hepatol 13, 324–337.

Kedinger, M., Duluc, I., Fritsch, C., Lorentz, O., Plateroti, M., and Freund, J.N. (1998). Intestinal epithelial-mesenchymal cell interactions. Ann N Y Acad Sci 859, 1–17.

Kretzschmar, K., and Clevers, H. (2017). Wnt/beta-catenin signaling in adult mammalian epithelial stem cells. Dev Biol 428, 273–282.

Li, Q., Sun, Y., Jarugumilli, G.K., Liu, S., Dang, K., Cotton, J.L., Xiol, J., Chan, P.Y., DeRan, M., Ma, L., et al. (2020). Lats1/2 Sustain Intestinal Stem Cells and Wnt Activation through TEAD-Dependent and Independent Transcription. Cell Stem Cell 26, 675–692 e678.

Ma, S., Meng, Z., Chen, R., and Guan, K.L. (2019). The Hippo Pathway: Biology and Pathophysiology. Annu Rev Biochem 88, 577–604.

Mah, A.T., Yan, K.S., and Kuo, C.J. (2016). Wnt pathway regulation of intestinal stem cells. J Physiol 594, 4837–4847.

McCarthy, N., Kraiczy, J., and Shivdasani, R.A. (2020). Cellular and molecular architecture of the intestinal stem cell niche. Nat Cell Biol 22, 1033–1041.

Muzumdar, M.D., Tasic, B., Miyamichi, K., Li, L., and Luo, L. (2007). A global double-fluorescent Cre reporter mouse. Genesis 45, 593–605.

Nusse, R., and Clevers, H. (2017). Wnt/beta-Catenin Signaling, Disease, and Emerging Therapeutic Modalities. Cell 169, 985–999.

Schindelin, J., Arganda-Carreras, I., Frise, E., Kaynig, V., Longair, M., Pietzsch, T., Preibisch, S., Rueden, C., Saalfeld, S., Schmid, B., et al. (2012). Fiji: an open-source platform for biological-image analysis. Nat Methods 9, 676–682.

Shoshkes-Carmel, M., Wang, Y.J., Wangensteen, K.J., Toth, B., Kondo, A., Massasa, E.E., Itzkovitz, S., and Kaestner, K.H. (2018). Subepithelial telocytes are an important source of Wnts that supports intestinal crypts. Nature 557, 242–246.

Valenta, T., Degirmenci, B., Moor, A.E., Herr, P., Zimmerli, D., Moor, M.B., Hausmann, G., Cantu, C., Aguet, M., and Basler, K. (2016). Wnt Ligands Secreted by Subepithelial Mesenchymal Cells Are Essential for the Survival of Intestinal Stem Cells and Gut Homeostasis. Cell Rep 15, 911–918.

Wirth, A., Benyo, Z., Lukasova, M., Leutgeb, B., Wettschureck, N., Gorbey, S., Orsy, P., Horvath, B., Maser-Gluth, C., Greiner, E., et al. (2008). G12-G13-LARG-mediated signaling in vascular smooth muscle is required for salt-induced hypertension. Nat Med 14, 64–68.

Yi, J., Lu, L., Yanger, K., Wang, W., Sohn, B.H., Stanger, B.Z., Zhang, M., Martin, J.F., Ajani, J.A., Chen, J., et al. (2016). Large tumor suppressor homologs 1 and 2 regulate mouse liver progenitor cell proliferation and maturation through antagonism of the coactivators YAP and TAZ. Hepatology 64, 1757–1772.

Yui, S., Azzolin, L., Maimets, M., Pedersen, M.T., Fordham, R.P., Hansen, S.L., Larsen, H.L., Guiu, J., Alves, M.R.P., Rundsten, C.F., et al. (2018). YAP/TAZ-Dependent Reprogramming of Colonic Epithelium Links ECM Remodeling to Tissue Regeneration. Cell Stem Cell 22, 35–49 e37.

Zanconato, F., Cordenonsi, M., and Piccolo, S. (2016). YAP/TAZ at the Roots of Cancer. Cancer Cell 29, 783–803.

Zhao, B., Wei, X., Li, W., Udan, R.S., Yang, Q., Kim, J., Xie, J., Ikenoue, T., Yu, J., Li, L., et al. (2007). Inactivation of YAP oncoprotein by the Hippo pathway is involved in cell contact inhibition and tissue growth control. Genes Dev 21, 2747–2761.

Zheng, Y., and Pan, D. (2019). The Hippo Signaling Pathway in Development and Disease. Dev Cell 50, 264–282.

